# DeepADR: Multi-modal Prediction of Adverse Drug Reaction Frequency by Integrating Early-Stage Drug Discovery Information via Kolmogorov-Arnold Networks

**DOI:** 10.1101/2025.07.29.667409

**Authors:** Jingting Wan, Chenyang Jia, Danhong Dong, Yigang Chen, Yang-Chi-Dung Lin, Yisheng He, Hsi-Yuan Huang, Hsien-Da Huang

**Affiliations:** School of Medicine, The Chinese University of Hong Kong, Shenzhen, Guangdong, 518172, P.R. China; Warshel Institute for Computational Biology, School of Medicine, The Chinese University of Hong Kong, Shenzhen, Guangdong, 518172, P.R. China; Guangdong Provincial Key Laboratory of Digital Biology and Drug Development, The Chinese University of Hong Kong, Shenzhen, Guangdong, 518172, P.R. China; Department of Endocrinology, Key Laboratory of Endocrinology of National Health Commission, Peking Union Medical College Hospital, Chinese Academy of Medical Sciences & Peking Union Medical College, Beijing, 100730, P.R. China

**Author notes:** To whom correspondence should be addressed. Yisheng He; Hsi-Yuan Huang; Hsien-Da Huang.

**Keywords:** adverse drug reactions, drug development, multi-modal deep learning, molecular representation, Kolmogorov–Arnold Network

## Abstract

Adverse drug reactions (ADRs) are a major cause of clinical trial failure and post-market withdrawal, posing significant risks to public health and impeding drug development. While computational methods offer an alternative to costly preclinical testing, existing models often fail with novel compounds by requiring pre-existing information such as drug-ADR associations or by inadequately integrating diverse data sources. Here, we introduce DeepADR, a multi-modal deep learning framework for predicting both the occurrence and frequency of ADRs using early-stage, readily available data. DeepADR integrates chemical structures and biological target profiles with semantic representations of ADR terms derived from a large language model. These heterogeneous parameters are fused using a Kolmogorov–Arnold Network (KAN), which effectively models complex, non-linear cross-modal interactions to capture underlying toxicological mechanisms. Our model outperforms existing methods in predicting both ADR occurrence and frequency, demonstrating robust generalization to new chemical entities. By effectively integrating chemical, biological, and semantic datasets, DeepADR provides a powerful, scalable tool for the early-stage safety assessment and candidate prioritization. This framework not only facilitates the prioritization of safer drug candidates but also offers a methodology for predicting the toxicity of other hazardous materials, holding significant promise for advancing public health.

## 1. Introduction

Ensuring drug safety throughout the pharmaceutical development pipeline is critical, as adverse drug reactions (ADRs) remain a substantial challenge. They contribute significantly to late-stage clinical trial failures, costly market withdrawals, and significant patient morbidity and mortality ^1,2^. Estimates indicate that ADRs are responsible for approximately 30% of drug development attrition and are among the leading causes of hospitalization in developed countries ^3^. Beyond the immediate threat to public health, these safety concerns impose significant financial burdens on the healthcare system and pharmaceutical industry. Traditional preclinical toxicological assessments form the foundation of safety evaluation. These include in vitro assays and animal testing. However, these experimental approaches have limitations. They involve high costs, lengthy timelines, restricted throughput, and challenges in accurately extrapolating findings to human physiology. ^4-6^ Consequently, there is a recognized need for effective computational methods. Such methods should be capable of predicting ADR risks early in the drug discovery process. This would improve decision making and help prioritize candidates with favorable safety profiles ^7^.

Computational ADR prediction has emerged as an important in silico approach. It complements traditional safety testing, particularly in the early stages where experimental data is scarce. These methods can be broadly categorized into drug-ADR association prediction and frequency prediction. Early efforts focused primarily on association prediction. The aim was to determine the likelihood of a link between a specific drug and a particular ADR. This was often framed as a binary or multi-label classification problem. Early approaches included Quantitative Structure-Activity Relationship (QSAR) models. These models correlate chemical structures or fragments with adverse reactions ^8^. Machine learning algorithms were also applied to chemical, biological, and phenotypic features ^9-14^. Examples include ensemble methods ^11,15^, recommender systems ^15^, bipartite local models (BLM) ^16^, restricted Boltzmann machines (RBM) ^17^, and attention-based networks ^18-20^. Notably, MSDSE ^18^ integrated multi-scale features derived from the drugs including word embedding, structure and substructure based embeddings by hybrid approaches combining self-attention. With the development of network-based methods, some studies leveraged drug and ADR similarity networks ^21,22^, linear neighborhood similarity ^23^, analysis of drug-target interactions and protein domains ^24^, or more complex network/domain information like NDDSA ^25^. The underlying principle was that similar drugs might elicit similar ADRs. Techniques like using inverse similarity and reliable negative samples were also explored to improve prediction accuracy ^26^. In addition, MVDSA ^20^ constructed knowledge graphs by integrating multiple relationship semantics and local topologies from the aspects of drug similarity, ADR similarity, and existing drug-ADR association matrix. While association-based models play an important role in post-marketing surveillance and risk identification, they often lack the resolution required for estimating the likelihood or severity of ADRs. These challenges have motivated a shift toward frequency prediction, which offers a more quantitative and clinically informative measure of ADR risk.

Frequency prediction focuses on estimating how often a specific ADR occurs in a population exposed to a given drug. This involves estimating how often a specific ADR occurs in a population exposed to a drug, providing a more quantitative and clinically relevant measure of risk. A notable contribution was made by Galeano et al. ^27^, who constructed a benchmark dataset based on SIDER ^28^ and employed matrix decomposition to predict ADR frequencies categorized into five levels. However, this initial work relied solely on known drug-ADR association data and did not incorporate any biological information. Subsequent research aimed to employ richer feature sets and more sophisticated algorithms. Based on the contribution, NRFSE ^29^ leveraged multi-view data, including chemical structure, GO annotation, MedDRA term ^30^, along with the drug-ADR association similarity to compute neighborhood regularization and make predictions. Numerous machine-learning and deep-learning approaches have been investigated for ADR frequency prediction. Zhao et al. first introduced MGPred ^31^, which employs graph attention networks (GAT) to integrate multiple data views, including drug-ADR neighbourhoods, molecular fingerprints, and ADR word embeddings. The same group subsequently proposed SDPred ^32^, extending this framework by incorporating additional feature types and modelling them with convolutional neural networks (CNN) to achieve richer ADR frequency predictions. DSGAT ^33^ converted each drug’s molecular structure and its ADR-based drug–ADR similarity matrix into graph-like images, which are then processed by GAT. Other GNN-based methods include idse-HE employing hybrid embeddings ^34^, iADRGSE combining graph embeddings and self-attention ^19^, and PreciseADR using heterogeneous GNNs on patient-centric graphs ^35^. Additionally, some methods focused on analyzing patient diagnosis subgraphs ^36^. Transformer architectures were also developed. HSTrans ^37^ learned unified embeddings from substructures. CrossFeat ^38^ used a CNN-transformer architecture with cross-attention for predicting frequencies for new drugs.

However, the features utilized by existing prediction tools are not well aligned with the type of information available during the early stages of drug discovery (Table 1). At this stage, the only accessible data typically include drug structures and drug targets ^39^. In contrast, information such as known ADRs of novel drugs and drug-related semantic features often becomes available only during clinical trials or after the drug has entered the market. Therefore, effectively leveraging drug structures, drug targets, and the semantic information of ADRs aligns most closely with the practical stages of drug development.

**Table 1.** Machine learning methods for identifying drug side effects based on SIDER.

| Method | Feature types for model training |  |  |  | Algorithms | Pub Med link |
| --- | --- | --- | --- | --- | --- | --- |
|  | molecular structure | drug target | ADR semantic information | drug-ADR associations or extensive information |  |  |
| Galeano’s (2019) |  |  |  | ✓ | matrix decomposition | 27 |
| MGPred (2021) | ✓ |  | ✓ | ✓ | GAT | 31 |
| SDPred (2022) | ✓ | ✓ | ✓ | ✓ | CNN | 32 |
| DSGAT (2022) | ✓ |  |  | ✓ | GAT | 33 |
| NRFSE (2023) | ✓ |  | ✓ | ✓ | matrix decomposition, neighborhood regularization | 29 |
| MSDSE (2024) | ✓ |  |  |  | self-attention network, CNN | 18 |
| CrossFeat (2024) | ✓ |  | ✓ |  | CNN, transformer, cross-attention | 38 |
| HSTrans (2025) | ✓ |  |  |  | transformer | 37 |
| DeepADR (our work) | ✓ | ✓ | ✓ |  | VAE, cross attention network, CNN, KAN | / |

Building on this rationale, we developed DeepADR (Figure 1a), a multimodal framework that integrates and enhances three distinct types of features for ADR association and frequency prediction. Molecular structures are represented by Molformer ^40^, a transformer architecture to learn chemically informed embeddings that capture local atomic topology, functional-group context, and long-range intramolecular interactions. Drug-target information is then encoded as multi-hot vectors and compressed with a Variational Autoencoder (VAE) ^41^ to capture nonlinear latent structure and reduce redundancy. Finally, semantic features of ADRs are extracted with BioBERT ^42^, a biomedical language model that enriches domain-specific terminology understanding. The structure and ADR semantic embeddings are jointly processed through a cross-attention mechanism followed by a CNN ^43,44^, which captures key cross-modal interactions and local hierarchical patterns. Finally, the integrated structural-semantic representations and the target embeddings are fused through a Kolmogorov–Arnold Network (KAN) ^45^, a recently proposed neural architecture that excels at modeling complex high-dimensional nonlinear relationships with strong approximation capacity. This design allows DeepADR to make accurate and data-efficient predictions of ADR associations and frequencies, supporting its translational relevance in preclinical safety evaluation by offering a proactive means to anticipate post-marketing safety concerns based on early-stage features in drug discovery.

**Figure 1.**
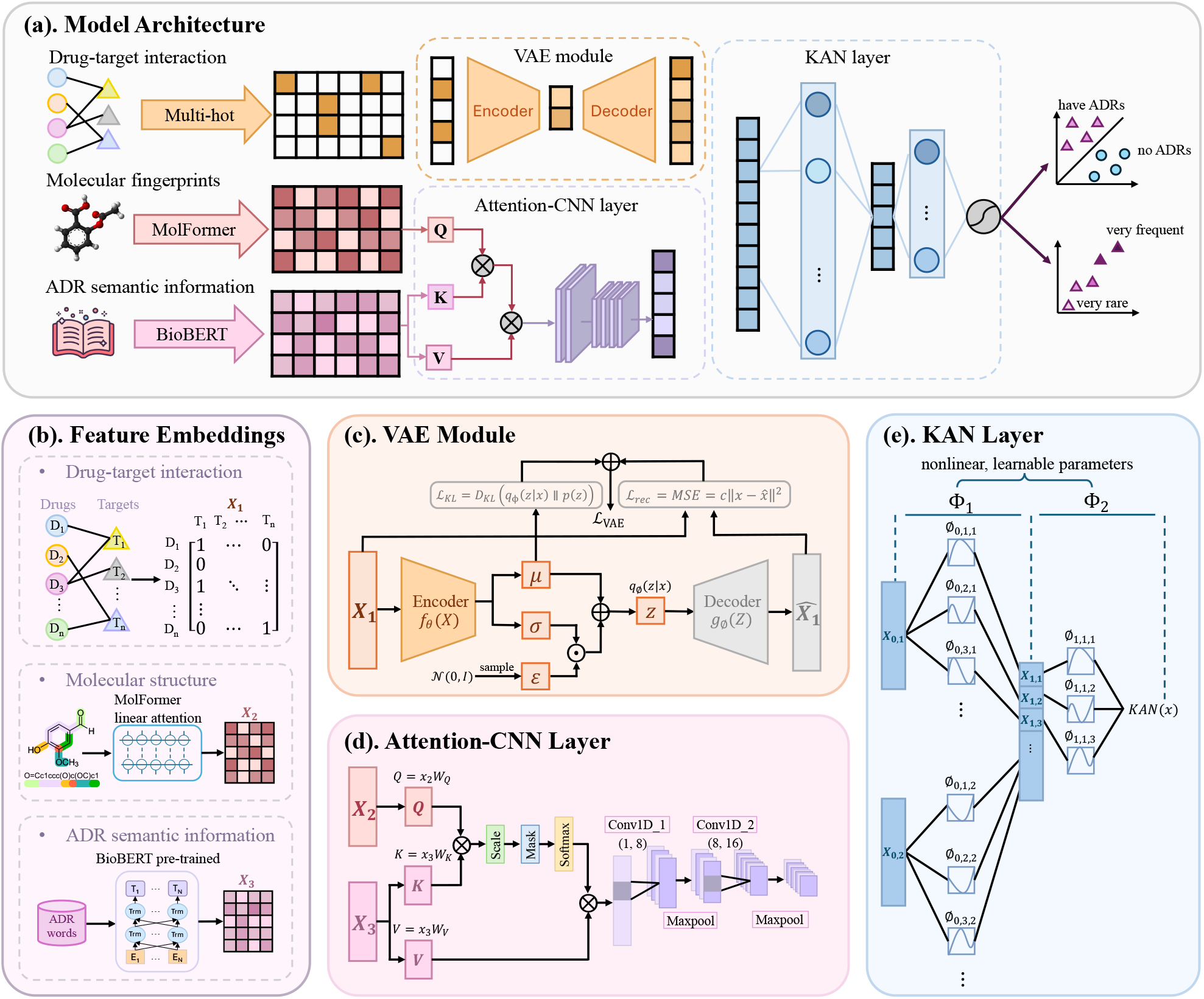
Model architecture and modular components of DeepADR. (a) Model architecture of the DeepADR framework. The model integrates drug–target profiles, molecular structure features, and ADR semantic information for multi-label frequency prediction. (b) Feature embedding process. Drug–target interactions are encoded as multi-hot vectors representing known associations, molecular fingerprints are extracted using MolFormer to capture structural properties, and ADR terms are embedded via a pretrained BioBERT model to obtain semantic representations. (c) VAE module. The multi-hot target vector is encoded into a latent distribution through neural networks estimating the mean and variance, sampled via the reparameterization trick, and decoded during training. The loss function consists of a binary cross-entropy reconstruction term and a KL divergence regularization term. (d) Attention-CNN module. This module enables interaction across modalities through attention mechanisms, followed by convolutional layers that capture localized dependencies within and between feature types. (e) KAN layer. The fused representation is passed through polynomial-based activation units, allowing the model to learn complex nonlinear interactions for precise ADR frequency prediction.

## 2. Materials and Methods

### 2.1. Data collection and preprocessing

To construct a comprehensive dataset for ADR prediction, we integrated drug-related and ADR-related information from multiple public databases (Figure 2, Supplementary Table S1). For ADR annotations, we utilized modified Galeano’s ^27^ benchmark dataset based on SIDER 4.1 ^28^, which contains curated drug–ADR associations with frequency-level annotations. These frequencies were categorized into five standard levels: Very rare (0 < *p* < 0.01 %), Rare (0.01 % ≤ *p* < 0.1 %), Infrequent (0.1 % ≤ *p* < 1 %), Frequent (1 % ≤ *p* < 10 %), and Very frequent (*p* ≥ 10 %), where *p* is the observed frequency of the ADR in the treated population. Molecular structures of drugs were retrieved from PubChem ^46^ in the form of SMILES strings, providing a basis for chemical feature extraction. Drug–target interaction data were obtained from DrugBank ^47^, enabling the incorporation of pharmacological profiles. To further standardize and group ADR terms for analysis, we mapped them to higher-level system organ classes using the MedDRA ontology. The final dataset thus combines structural, pharmacological, and semantic information, supporting the development of a multi-modal predictive model. The final dataset contains 719 drugs, 994 ADRs, and a total of 33,157 non-zero entries representing observed drug–ADR pairs. An equal number of randomly sampled negative pairs were used to construct the dataset for binary classification. All positive pairs were retained for the frequency regression task. For performance evaluation, we split the dataset into training, validation, and test sets with the portion of 64%, 16% and 20% respectively. Model development and hyperparameter tuning were performed using five-fold cross-validation.

**Figure 2.**
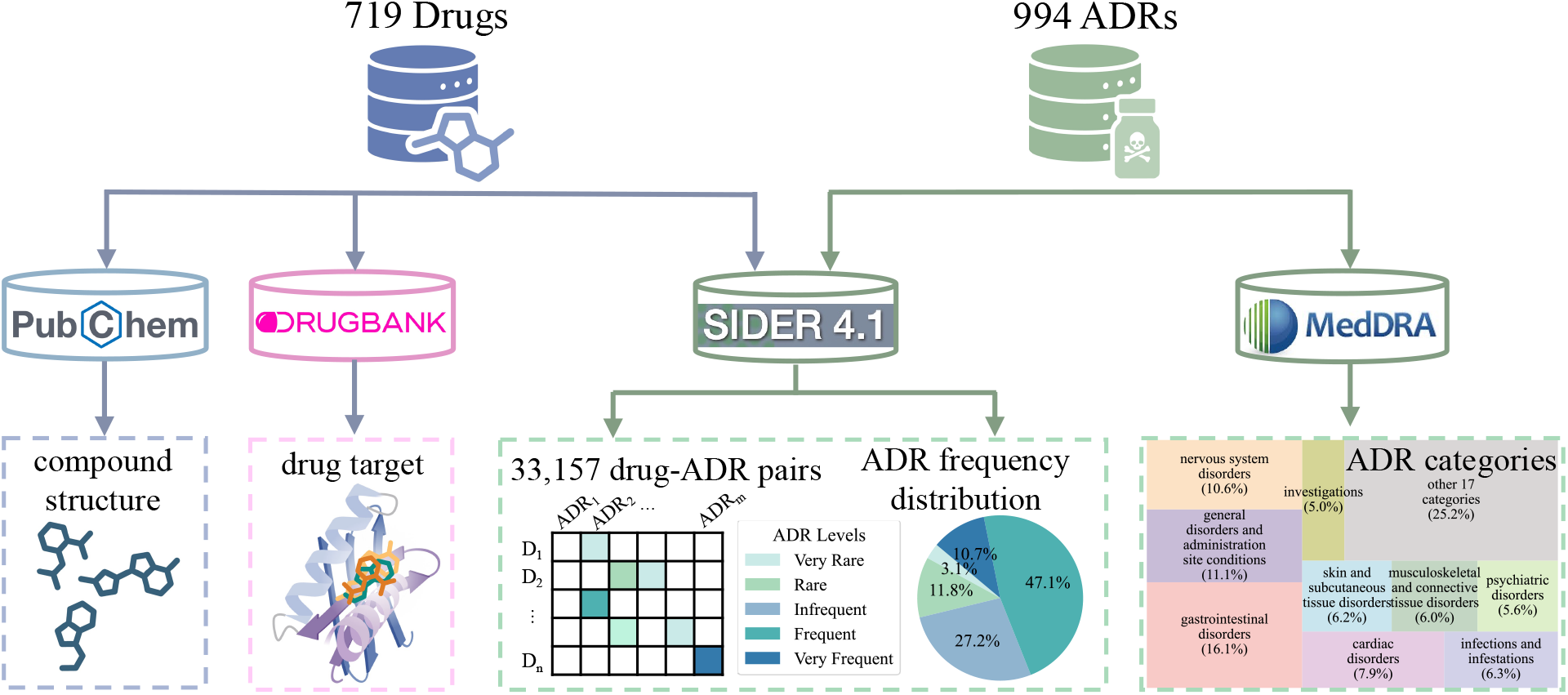
Integrated data sources and extracted features for ADR analysis. Chemical structures and target profiles were retrieved from PubChem and DrugBank. ADR frequency data were obtained from SIDER 4.1, while standardized ADR categories were mapped using MedDRA for analysis.

### 2.2. Feature embedding

Three modalities of input features were embedded for each drug–ADR pair (Figure 1b).

#### 2.2.1. Multi-hot encoding for drug-target interaction embedding

Each drug’s target profile is represented by a multi-hot vector *t ∈* {0,1}^*d*^, where *d* is the total number of possible protein targets. For a given drug, *t*_*j*_ = 1 if the drug is known to interact with target *j*, and *t*_*j*_ = 0 otherwise. This encoding captures the binary interaction status across all targets and forms the input to the model training process.

#### 2.2.2. Molformer for molecular embedding

To extract structural representations from SMILES strings, we employed Molformer ^40^, a Transformer-based masked language model specifically designed for molecular sequences.

The input SMILES string is first tokenized into a sequence *x* = {*x*_1_, *x*_2_, …, *x*_*L*_}, where each token *x*_*i*_ mapped to an embedding vector ***e***_*i*_ *∈* ℝ^*d*^. These token embeddings are combined with absolute positional encodings ***p***_*i*_ *∈* ℝ^*d*^ to form the input to the encoder:

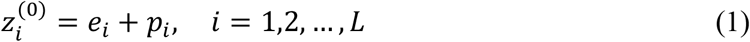

Molformer then applies a stack of multi-head self-attention and feed-forward layers to model the contextual dependencies between molecular tokens. The self-attention mechanism computes attention scores between each query–key pair via inner products, followed by a softmax-based aggregation. Specifically, the attention output for the *m*-th token is given by:

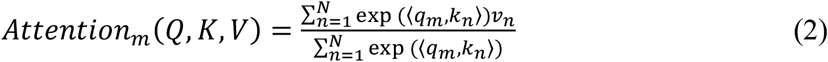

where Q, K and V are the query, key and value respectively.

#### 2.2.3. BioBERT for ADR semantic embedding

To capture the semantic meaning of ADR terms, we used BioBERT ^42^, a pre-trained biomedical language model based on the BERT ^48^ architecture and trained on large-scale biomedical corpora. Each ADR term was treated as a short text sequence and tokenized using WordPiece tokenization. The resulting token sequence ***s*** = {*s*_1_, *s*_2_, …, *s*_*M*_} was passed through BioBERT to obtain contextualized token embeddings {*h, h*_2_, …, *h*_*M*_} *∈* ℝ^*d*^. To derive a fixed-length semantic representation for each ADR, we used the output embedding of the [CLS] token, which serves as a global summary of the input sequence:

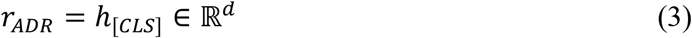

This vector encodes the semantic and contextual information of the ADR term in the biomedical domain. The resulting embedding was used as the input representation for the ADR modality in the subsequent multi-modal prediction framework.

### 2.3. Multi-modal prediction

To model the complex relationships between drugs and ADRs, DeepADR integrates molecular structure features, ADR semantic embeddings, and pharmacological target representations in a unified multi-modal deep learning framework (Figure 1c-e).

#### 2.3.1. Variational Autoencoder (VAE)

The multi-hot target vector *t* is encoded into latent space using a Variational Autoencoder ^41^. The encoder produces parameters of a Gaussian distribution:

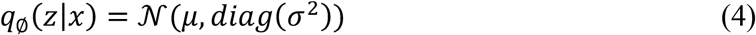

where μ = *f*_C_1*x*2 and σ = *f*_D_1*x*2 are learned through neural networks. The decoder reconstructs an approximation of the original multi-hot vector from latent *z*, denoted as 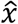:

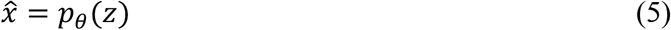

The VAE loss function combines reconstruction loss and KL divergence:

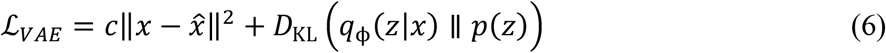

where *p*1*z*2 is a standard Gaussian prior.

#### 2.3.2. Attention mechanism

To model the interaction between drug structure and adverse reaction semantics, we apply a cross-attention mechanism in which molecular embeddings act as queries, while semantic ADR embeddings act as keys and values ^43^. Specifically, let the molecular embedding from Molformer be 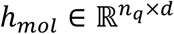, and the ADR embedding from BioBERT be 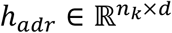, where *d* is the hidden dimension and *n*_6_, *n*_7_ are the number of tokens (or pooled representations) for molecule and ADR, respectively.

The cross-attention is computed as:

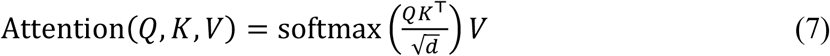

where:

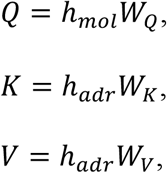

and *W*_*Q*_, *W*_*K*_, *W*_*V*_ *∈* ℝ^*d*×*d*^ are trainable projection matrices. The output of this attention mechanism is denoted as *H*_*attn*_, which serves as the input to the convolutional layer.

#### 2.3.3. CNN layer

A two-layer 1D CNN ^44^ is employed to extract local and hierarchical features from the cross-attention output *H*_*attn*_. The computation proceeds as follows:

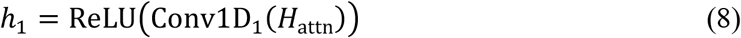

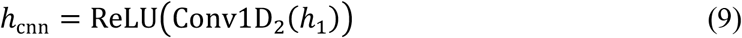

This two-stage convolutional block captures hierarchical feature interactions between molecular and ADR representations. The intermediate output *h*_1_ represents the locally filtered activation map produced by the first convolutional layer. The second layer further refines this representation to produce the final structural-semantic embedding *h*_cnn_, which serves as input to the subsequent fusion module.

#### 2.3.4. KAN layer

To capture complex nonlinear relationships between the structural-semantic representation and pharmacological target embeddings, we adopt KAN for final feature fusion and prediction ^45^. The KAN model is built upon the Kolmogorov–Arnold representation theorem, which states that any continuous multivariate function can be decomposed into a finite sum of compositions of univariate functions.

Formally, let *x* = [*h*_*CNN*_; *z*] *∈* ℝ^*k*^ be the concatenated feature vector from the CNN and VAE branches. The KAN output is given by:

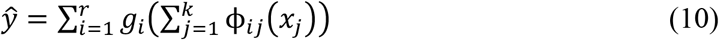

where:

ϕ_*ij*_ are trainable univariate basis functions,

*g*_*i*_ are learnable outer functions,

*r* is the number of summation terms,

ŷ is the predicted ADR frequency.

### 2.4. Model Evaluation

We evaluated the model performance under the following methods across both classification and regression tasks.

#### 2.4.1. Area Under the Receiver Operating Characteristic curve (AUROC)

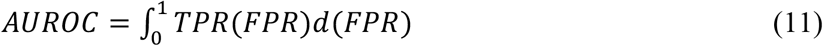

where:

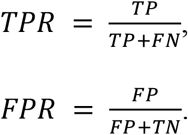

#### 2.4.2. Area Under the Precision-Recall curve (AUPR)

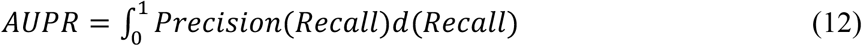

where:

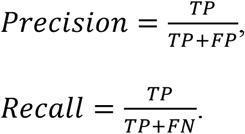

#### 2.4.3. F1-score

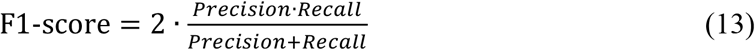

#### 2.4.4. Root Mean Squared Error (RMSE)

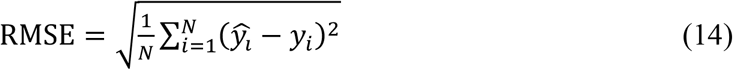

where 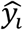 is the predicted value, *y*_*i*_ is the ground truth, and *N* is the number of samples.

#### 2.4.5. Mean Absolute Error (MAE)

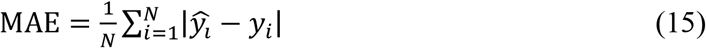

#### 2.4.6. Pearson Correlation Coefficient (PCC)

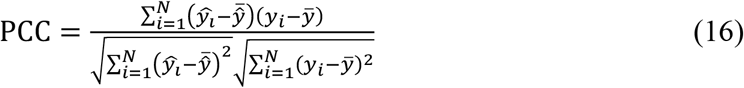

where:

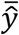 is the average of predicted values,

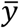 is the average of the ground truth values

## 3. Results

### 3.1. Performance Comparison Across Classification and Regression Tasks

We performed a comprehensive comparative analysis of DeepADR against seven state-of-the-art models, including Galeano’s ^27^, MGPred ^31^, SDPred ^32^, DSGAT ^33^, NRFSE ^29^, MSDSE ^29^ and HSTrans ^37^, using five independent repeated tests with distinct test sets (Table 2). All models were strictly constrained to utilize only information accessible at the early stages of drug discovery. In the classification task, DeepADR achieved the highest overall performance, demonstrating superior discrimination ability with an AUROC of 0.9346, an AUPR of 0.9305, and an F1 score of 0.8618. While DSGAT ^33^ and NRFSE ^29^ also exhibited relatively strong classification performance (AUROC > 0.91, F1 score > 0.82), their regression outcomes were substantially weaker, suggesting these methods primarily excel in binary association prediction rather than frequency estimation. MGPred ^31^ and SDPred ^32^ presented moderate classification capabilities (AUROC < 0.80), indicating limited accuracy in predicting ADR occurrence.

**Table 2.**
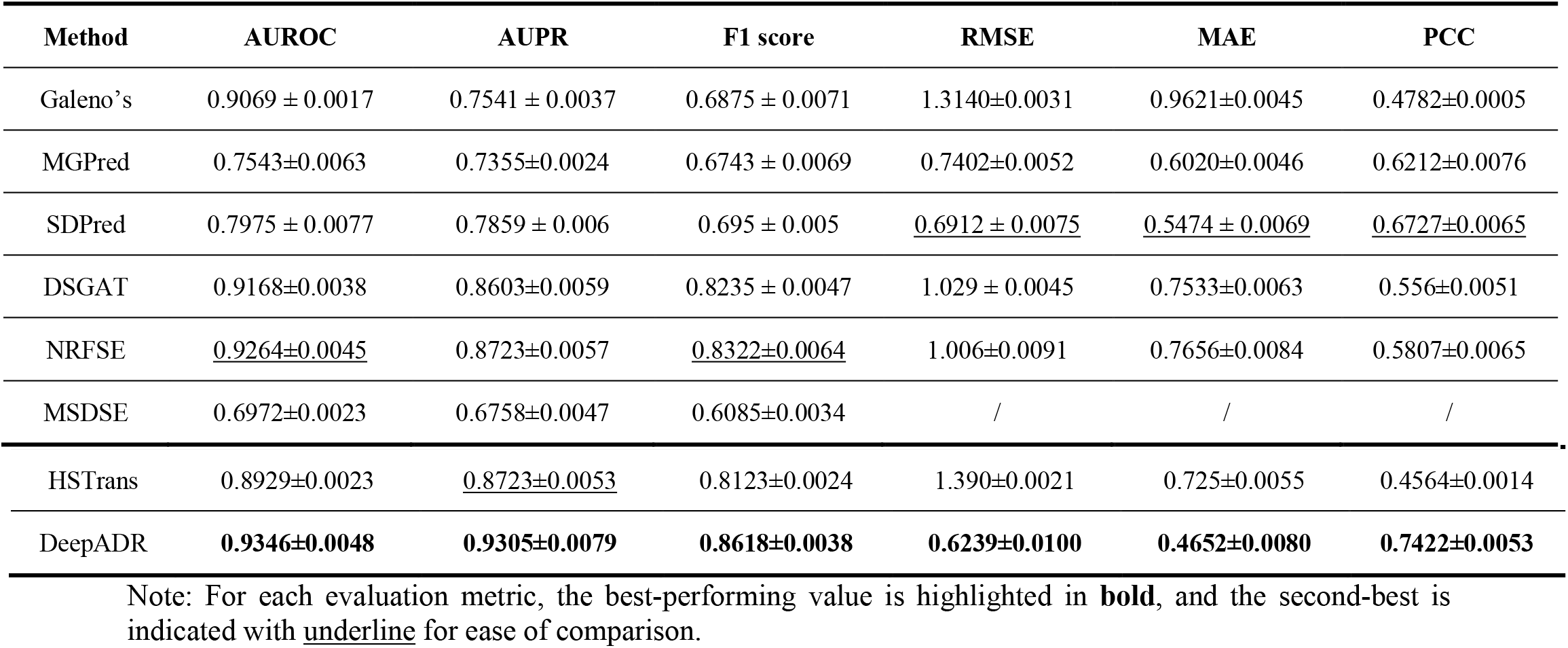
Performance comparison of ADR frequencies between DeepADR and other models.

Regarding regression, DeepADR significantly outperformed all other models by achieving the lowest RMSE (0.6239), lowest MAE (0.4652), and the highest PCC (0.7422). The simultaneously low RMSE and MAE values indicate that DeepADR predictions are both precise and consistent, demonstrating robustness to potential outliers in the data. Meanwhile, the high PCC reflects a strong correlation between predicted and actual ADR frequencies, confirming the model’s accuracy in capturing quantitative relationships even in the presence of variability or outliers. In contrast, Galeano’s approach ^27^ and HSTrans ^37^ showed inferior regression performance (RMSE > 1.3), reflecting their limited capacity to quantify ADR frequency accurately without adequately using accessible information in drug discovery.

These results underscore the efficacy and robustness of DeepADR’s multimodal integration strategy, emphasizing its unique capability to maintain high performance in both classification and regression tasks when data is restricted to early-stage drug discovery information. Consequently, DeepADR offers substantial improvements for early and accurate ADR risk prediction, facilitating more informed and safer drug candidate prioritization in preclinical phases.

### 3.2. Ablation Studies of DeepADR on Features and Algorithm Modules

We evaluated the contribution of individual features and architectural components to the predictive performance of DeepADR using six distinct metrics, including AUROC, AUPR, and F1 score for classification tasks, as well as RMSE, MAE, and PCC for regression tasks (Figure 3, Supplementary Table S2). It is shown that any removal of any of the three early-stage drug feature modalities, including chemical structure, biological targets, or ADR semantics, results in performance degradation across all metrics (Figure 3a). Among these, omitting ADR semantic embeddings leads to the most substantial decline, as they constitute the sole source of information on the ADR side and account for half of the predictive input space. These findings highlight that effective safety prediction requires integrating complementary chemical, biological, and semantic information.

**Figure 3.**
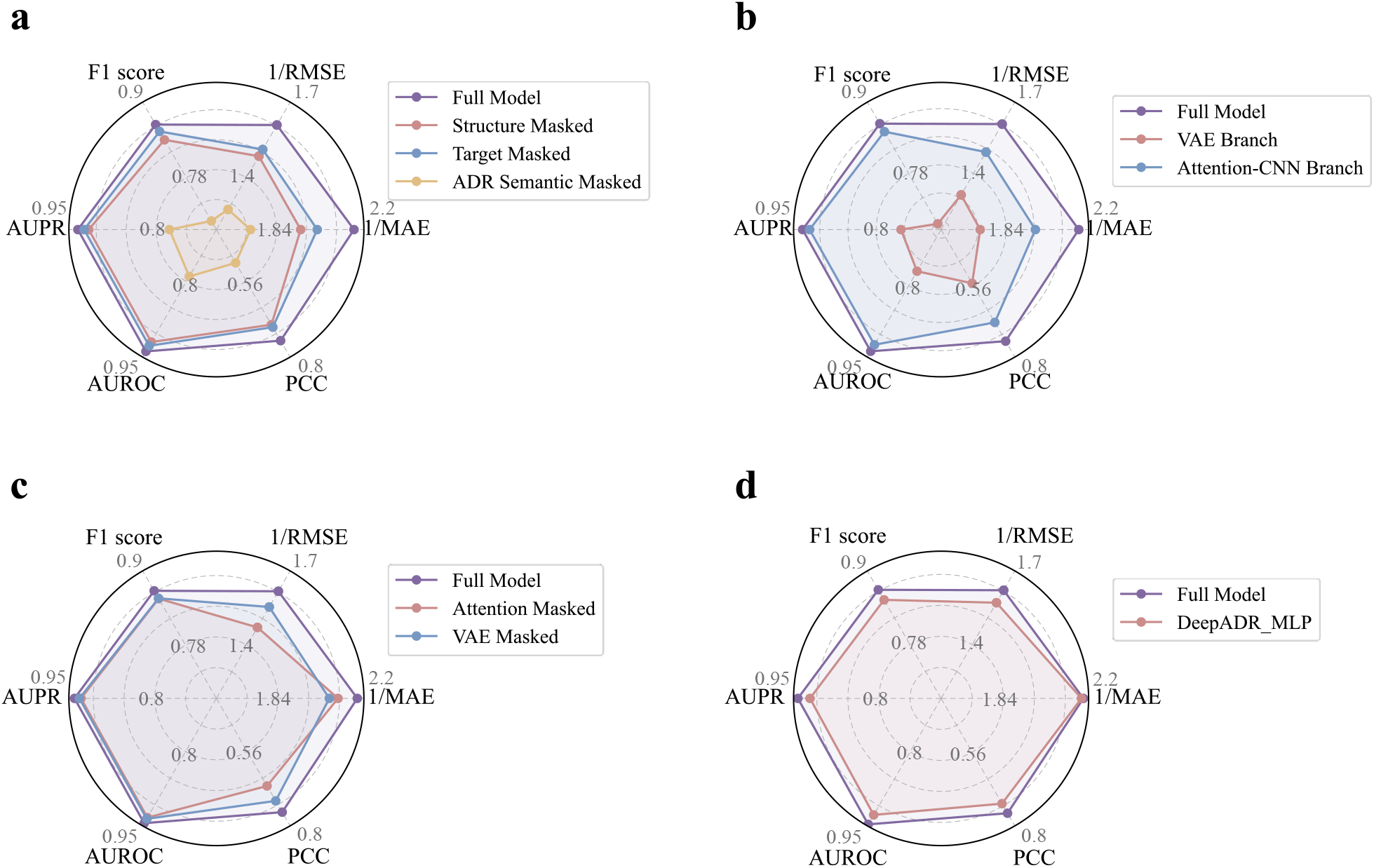
Ablation studies of the DeepADR model. Radar plots present six evaluation metrics (AUROC, AUPR, F1 score, 1/RMSE, 1/MAE, and PCC); larger areas indicate better performance. (a) Feature-mask experiment in which structure, target, or ADR-semantic inputs are individually removed. (b) Branch-only experiment that isolates the VAE branch or the Attention CNN branch. (c) Module-mask experiment that omits the cross-attention block or the VAE encoder. (d) Fusion-layer experiment that replaces the KAN with MLP.

Further ablation studies revealed specialized roles for distinct branches within the network (Figure 3b-d). Specifically, retaining only the Attention-CNN branch, responsible for integrating chemical structure and ADR semantics, largely preserves classification capabilities but adversely affects frequency estimation accuracy. Conversely, isolating the VAE branch, which generates compact pharmacological representations from target profiles, results in a marked decline across all metrics, with a more significant impact on classification performance (Figure 3b). These results confirm that the joint contribution of both branches is critical to the overall balanced performance of the complete model. Module-level ablations provided additional insights into component functionality (Figure 3c). Disabling the cross-attention module weakens the integration between molecular and semantic modalities, while removal of the VAE encoder impairs the representational richness of target features. In each scenario, performance metrics decline toward intermediate levels between single-branch models and the full architecture. Additionally, replacing the KAN with a multiple lay perceptron (MLP) module consistently reduces all evaluation metrics, indicating that KAN more effectively models the high-dimensional, nonlinear relationships essential for simultaneous optimization of classification and regression objectives (Figure 3d, Supplementary Figure S1a-b). Moreover, during training, the validation loss of KAN declines more rapidly and smoothly than that of the MLP, suggesting that KAN not only facilitates more stable convergence but also achieves better generalization (Supplementary Figure S1c). This training behavior reflects KAN’s capacity to extract essential cross-domain biological signals, supporting improved robustness in early-stage ADR forecasting. Collectively, these ablation analyses demonstrate that each early-stage modality and each architectural component provides measurable and indispensable contributions to the robust predictive performance of DeepADR.

### 3.3. Performance Across MedDRA-Classified ADR Categories

To evaluate the robustness of DeepADR across diverse physiological systems, we analyzed its regression performance by ADR category using high-level group terms from the MedDRA ontology. As shown in Figure 4, DeepADR maintains consistently low mean absolute error (MAE) across most categories, highlighting its strong generalization ability (Supplementary Table S3). Categories with abundant training data, such as *infections and infestations, general disorders and administration site conditions*, and *gastrointestinal disorders*, stably exhibit particularly high prediction accuracy. In contrast, categories with limited data, such as *pregnancy, puerperium and perinatal* conditions, *congenital, familial and genetic disorders* and *surgical and medical procedures*, show relatively more variable MAE. This variability may be attributed to the skewed label distribution common in low-sample scenarios, which can lead to either very high or deceptively low MAE depending on frequency clustering. Additionally, categories like *immune system disorders* exhibit slightly elevated MAE, potentially due to the underlying mechanistic heterogeneity. Overall, DeepADR demonstrates strong and consistent performance across a broad spectrum of ADR types.

**Figure 4.**
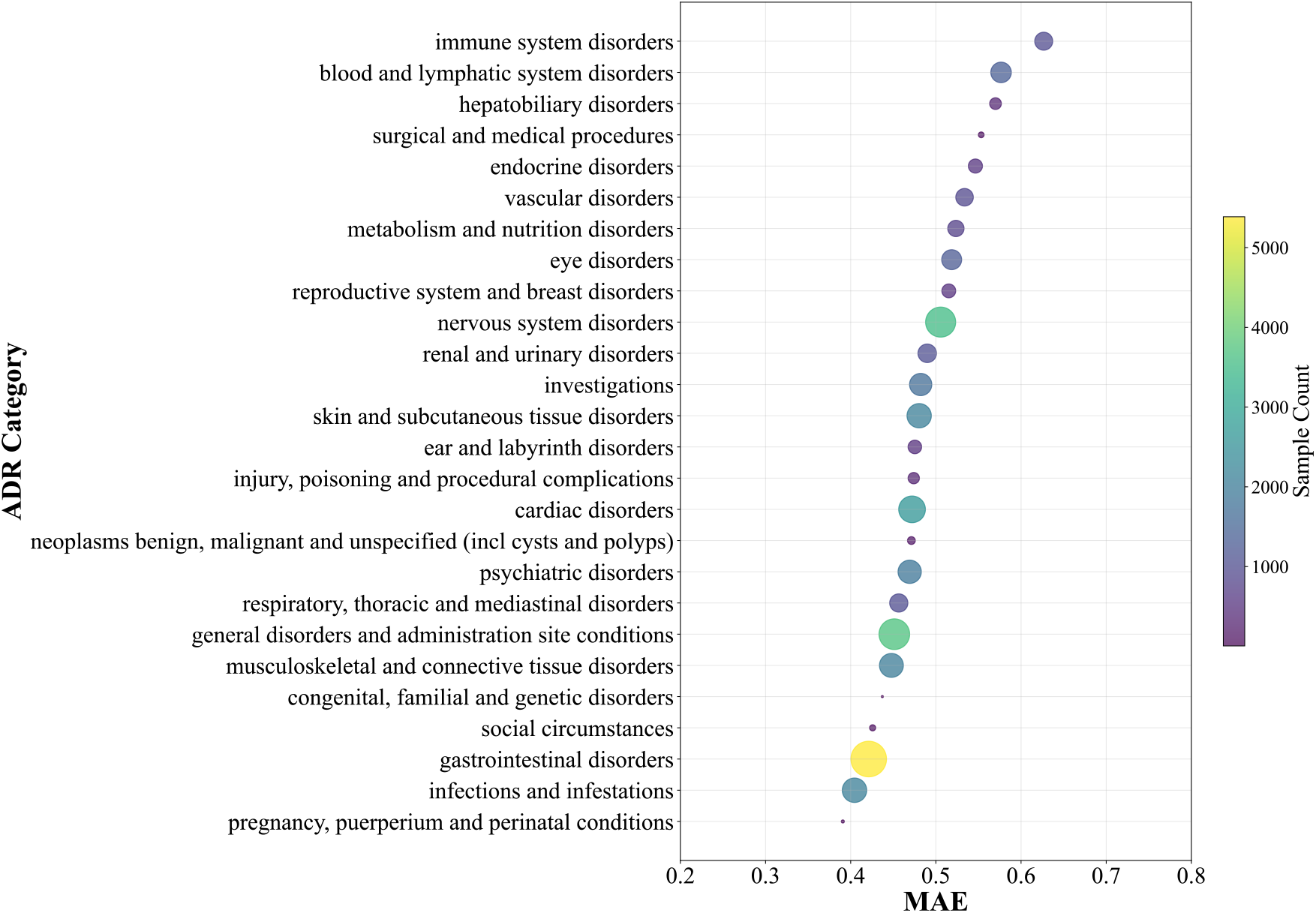
ADR category-wise prediction performance based on MAE. Each point represents a MedDRA-defined high-level ADR category. The x-axis indicates the MAE of DeepADR predictions within that category, while bubble size and color correspond to the number of samples (log-scaled). Categories with fewer samples tend to exhibit higher variability in prediction accuracy.

### 3.4. Case Studies of DeepADR Model

To further illustrate the practical applicability and predictive robustness of DeepADR, we conducted detailed case studies from both drug-centric and ADR-centric perspectives (Figure 5a-b, Supplementary Table S4-5). For the drug-centric evaluation, we focused on *Carbamazepine*, an anticonvulsant widely used in epilepsy and bipolar disorder ^49,50^, and *Terbinafine*, a commonly prescribed antifungal agent for dermatophyte infections ^51,52^. Both drugs have diverse ADR profiles spanning multiple frequency levels. DeepADR accurately identified frequent ADRs, as well as clinically significant rare ADRs such as agranulocytosis, anaphylactic shock, angioedema, and pancytopenia, emphasizing the model’s sensitivity in capturing rare safety risks.

**Figure 5.**
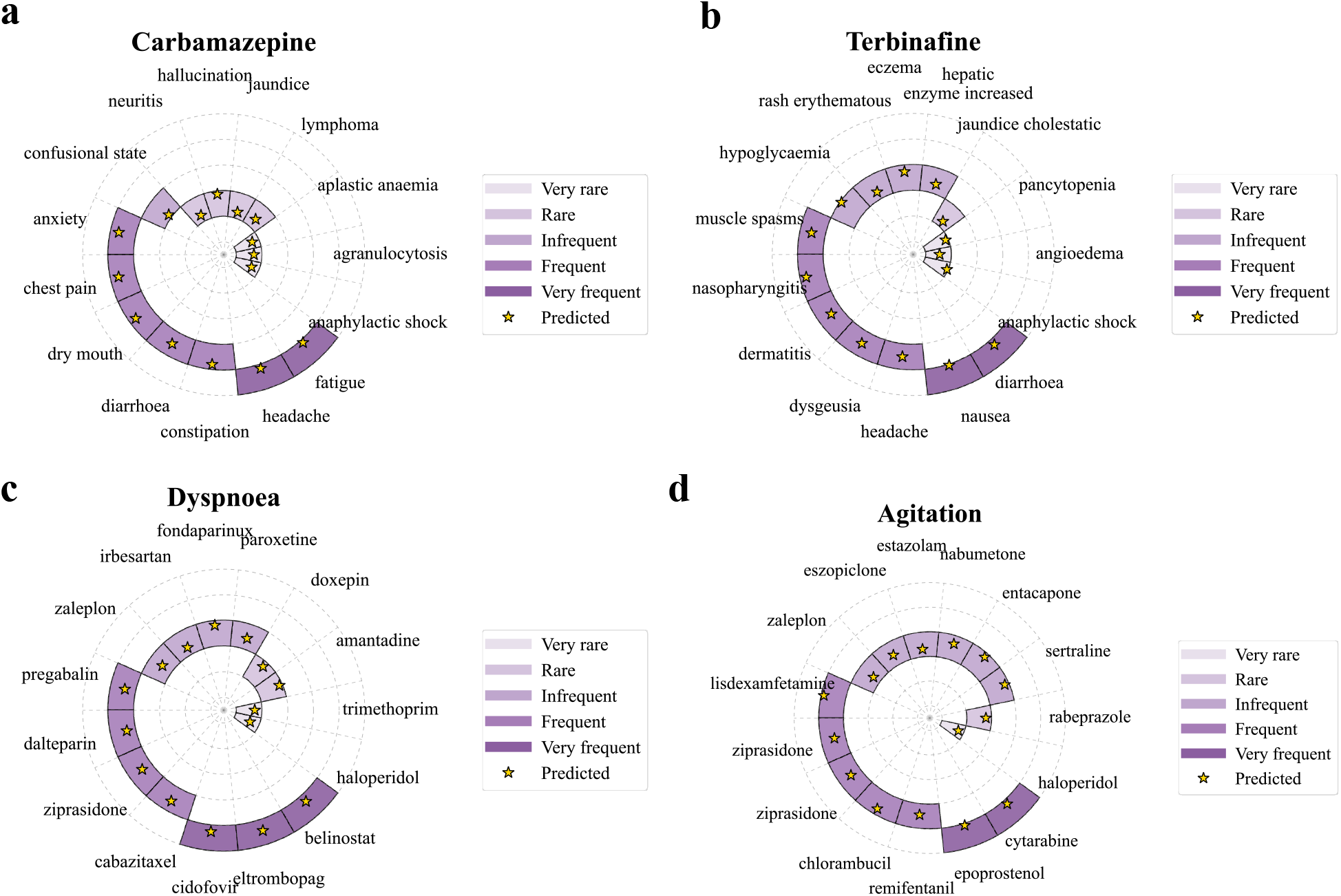
Case studies of ADR frequency prediction. Drug-centric case studies for (a) *Carbamazepine* and (b) *Terbinafine*, showing associated ADRs with color-coded frequency levels. ADR-centric case studies for (c) *Dyspnoea* and (d) *Agitation*, showing associated drugs and their reported frequencies. Stars indicate predictions made by DeepADR. Each prediction is produced as a continuous regression output and subsequently mapped to the nearest of the five frequency levels for visual comparison with reported data.

For the ADR-centric analysis, we examined *Dyspnoea*, a clinically important respiratory symptom ^53^, and *Agitation*, a common neuropsychiatric adverse reaction (Figure 5c-d, Supplementary Table S6-7) ^54^. DeepADR correctly predicted the frequency profiles for multiple drugs associated with these ADRs. For instance, *Dyspnoea* was accurately linked to drugs including pregabalin and ziprasidone, while *Agitation* was associated with drugs such as sertraline and haloperidol. The close alignment between model predictions and known ADR frequencies highlights DeepADR’s effectiveness in leveraging diverse chemical and pharmacological information for precise frequency estimation.

Collectively, these case studies demonstrate DeepADR’s proficiency in providing accurate, frequency-specific ADR predictions, highlighting its potential to enhance early-stage drug safety assessment and clinical risk management.

## 4. Discussion

The process of drug development typically unfolds across multiple stages, beginning with target identification and validation during the early discovery phase, where potential druggable proteins or pathways are determined through genomic, proteomic, or functional assays ^39,55^. This is followed by lead compound optimization, preclinical testing, and clinical trials, ultimately culminating in regulatory approval and post-marketing surveillance ^56^. While drug safety assessment is formally introduced during preclinical and clinical testing, much of the most comprehensive safety information, especially regarding rare or long-term ADRs, is only uncovered through post-marketing pharmacovigilance data ^1^. However, incorporating such late-stage information retrospectively into safety assessment pipelines is often too delayed to impact candidate selection or design modifications.

In addition, recent advances in large language models (LLMs) have facilitated the embedding of biomedical concepts into dense semantic spaces, enabling ADR terms to be represented with rich contextual and ontological information ^42,48^. Since MedDRA ADR terminology ^30^ is independent of any specific compound and remains stable regardless of the existence of a particular drug, these embeddings can be validly utilized as input features for learning tasks even in early-stage modeling. In contrast, the use of drug-side semantic information, which typically derives from literature mining, clinical narratives, or accumulated pharmacological knowledge, is far less feasible for newly designed compounds ^18^. This is because such representations inherently require large volumes of prior data ^42,57,58^ that are unavailable at the onset of drug development. From a translational standpoint, we therefore argue that the most practically accessible features in the early discovery stage are limited to chemical structure and known or predicted drug–target interactions. These data types are obtainable immediately after molecular design and prior to extensive experimental or clinical characterization, making them suitable for real-time safety evaluation frameworks.

By leveraging early-stage features to estimate post-marketing ADR frequencies, DeepADR helps bridge the gap between preclinical modeling and clinical safety outcomes. Unlike retrospective assessments, the model forecasts risk before any human exposure, enabling earlier risk stratification and proactive candidate prioritization. For example, within series that share a target or chemotype, DeepADR can rank parent compounds and derivatives by predicted occurrence and frequency, flag ADR categories that rise with specific substitutions or substructures, and identify safer analogues within equipotent sets, thereby focusing experimental effort on candidates with more favorable safety profiles.

Although the framework performs well overall, several limitations remain. First, model accuracy is lower for sparsely represented ADR categories, such as immune-system disorders, where limited data and complex biology pose challenges to representation learning. Second, external validation against real-world pharmacovigilance databases has not yet been undertaken. Future work will explore techniques for few-shot learning in rare ADR categories, and evaluate the model on independent clinical datasets to strengthen its generalizability.

## 5. Conclusion

This study introduces DeepADR, a multimodal framework that predicts both the presence and frequency of adverse drug reactions using only information accessible at the earliest stage of drug discovery, namely molecular structure, drug-target profiles, and standardized ADR terminology. Through nonlinear fusion implemented by a KAN, the model demonstrates strong predictive power and provides a practical tool for preclinical safety evaluation. By anticipating post-marketing ADR risks from preclinical data, DeepADR can inform early decision-making, improve compound prioritization, and reduce post-market failure. Future refinements will focus on improving performance in under-represented ADR categories and validating the model with real-world safety data.

## Data Availability

The drug–ADR associations were obtained from the SIDER database (http://sideeffects.embl.de/).

Molecular structures in SMILES format were retrieved from PubChem (https://pubchem.ncbi.nlm.nih.gov/), and drug target information was sourced from DrugBank (https://go.drugbank.com/).

## Code Availability

Source code used in this study are available at the GitHub repository: https://github.com/Wjt777/DeepADR.

## Acknowledgements

This research benefited significantly from the interdisciplinary research environment and cut-ting-edge instrumentation maintained by the Computational Platform of Warshel Institute for Computational Biology. Their ongoing commitment to research infrastructure development has been crucial to our scientific endeavors. Authors are also grateful to the library of The Chinese University of Hong Kong, Shenzhen for providing effective database service. We would like to express our sincere gratitude to the Vincent & Lily Woo Foundation for their generous support of the Vincent & Lily Woo Fellowship in Memory of Dr Albert Wong. This fellowship, endowed by the Vincent & Lily Woo Foundation, is provided through MCMIA Foundation Limited, and we are deeply grateful for their contribution to our research.

## Author Contributions

J.W. conceived the study, curated data, developed methodology, performed formal analyses, generated visualizations, and wrote the original draft. C.J. contributed to data curation, wrote parts of the original draft, and participated in review and editing. D.D. assisted with methodology, produced visualizations, and revised the manuscript. Y.C. contributed to methodology and manuscript editing. Y-C-D.L. reviewed and edited the manuscript. Y.H. provided methodological guidance and participated in review. H-Y.H. contributed to methodology development, supervised the work, and critically revised the manuscript. H-D.H. provided overall supervision, led conceptualization, offered methodological guidance, and approved the final revision. All authors read and approved the final manuscript.

