## Supplementary Information for "DeepADR: Multi-modal Prediction of Adverse Drug Reaction Frequency by Integrating Early-Stage Drug Discovery Information via Kolmogorov-Arnold Networks"

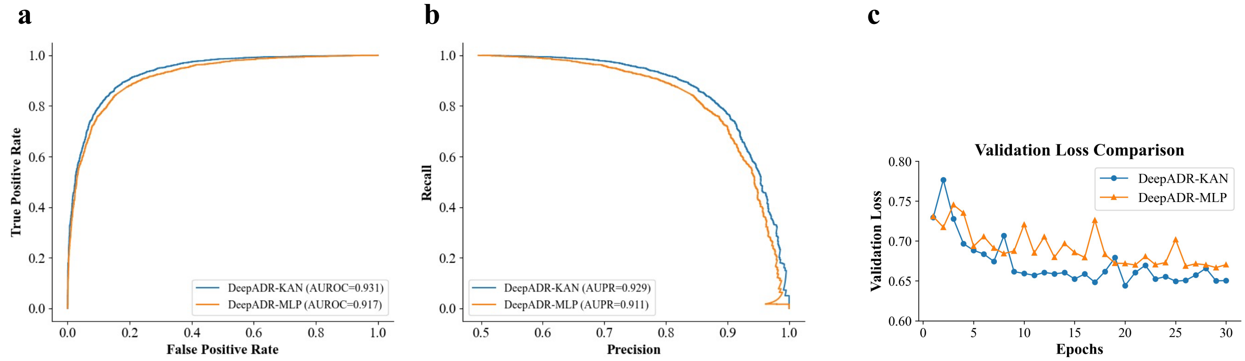


**Supplementary Figure S1. Comparison between KAN-based and MLP-based fusion in DeepADR.** (a) ROC and (b) PR curves for the classification task. (c) Validation-loss trajectories over 30 training epochs.

**Supplementary Table S1.** Details of Dataset

| Data Type | Data Number |
|---|---|
| **Drug** | 719 |
| **ADR** | 994 |
| **Positive Drug-ADR pairs** | 33,157 |
| **Zero-entries** | 681,529 |
| **High-level ADR categories** | 27 |

**Supplementary Table S2.** Ablation Study Results

| | | 1/RMSE | 1/MAE | PCC | AUROC | AUPR | F1 score |
|---|---|---|---|---|---|---|---|
| **Feature Masking** | Structure Masked | 1.4828 | 1.9369 | 0.693 | 0.9165 | 0.9125 | 0.8383 |
| | Target Masked | 1.5085 | 2.0028 | 0.7007 | 0.9238 | 0.9201 | 0.8512 |
| | ADR Semantic Masked | 1.2776 | 1.7364 | 0.5025 | 0.7903 | 0.7781 | 0.7133 |
| **Branch Removal** | VAE Branch | 1.3122 | 1.7301 | 0.5521 | 0.7703 | 0.756 | 0.6974 |
| | Attention-CNN Branch | 1.4879 | 1.9650 | 0.6806 | 0.9212 | 0.9181 | 0.8487 |
| **Module Masking** | Attention Masked | 1.4670 | 2.0746 | 0.6634 | 0.9241 | 0.9199 | 0.85 |
| | VAE Masked | 1.5438 | 2.0404 | 0.7083 | 0.9264 | 0.9225 | 0.8505 |
| **Fusion Method** | DeepADR_MLP | 1.5564 | 2.1422 | 0.7136 | 0.917 | 0.911 | 0.8468 |
| **Full Model** | | **1.6028** | **2.1496** | **0.7422** | **0.9346** | **0.9305** | **0.8618** |

**Supplementary Table S3.** ADR category-wise prediction performance

| ADR | Sample Count | RMSE | MAE |
|---|---|---|---|
| blood and lymphatic system disorders | 1,364 | 0.7414 | 0.5765 |
| cardiac disorders | 2,626 | 0.6408 | 0.4720 |
| congenital, familial and genetic disorders | 5 | 0.4931 | 0.4371 |
| ear and labyrinth disorders | 484 | 0.6065 | 0.4753 |
| endocrine disorders | 535 | 0.7206 | 0.5464 |
| eye disorders | 1,267 | 0.6783 | 0.5185 |
| gastrointestinal disorders | 5,387 | 0.5734 | 0.4211 |
| general disorders and administration site conditions | 3,717 | 0.6151 | 0.4512 |
| hepatobiliary disorders | 337 | 0.7498 | 0.5700 |
| immune system disorders | 979 | 0.8218 | 0.6266 |
| infections and infestations | 2,105 | 0.5653 | 0.4044 |
| injury, poisoning and procedural complications | 311 | 0.6217 | 0.4740 |
| investigations | 1,669 | 0.6729 | 0.4821 |
| metabolism and nutrition disorders | 765 | 0.6782 | 0.5235 |
| musculoskeletal and connective tissue disorders | 1,997 | 0.6257 | 0.4478 |
| neoplasms benign, malignant and unspecified (incl cysts and polyps) | 122 | 0.6292 | 0.4712 |
| nervous system disorders | 3,533 | 0.6691 | 0.5056 |
| pregnancy, puerperium and perinatal conditions | 12 | 0.4781 | 0.3907 |
| psychiatric disorders | 1,868 | 0.6371 | 0.4692 |
| renal and urinary disorders | 1,055 | 0.6599 | 0.4898 |
| reproductive system and breast disorders | 508 | 0.6779 | 0.5152 |
| respiratory, thoracic and mediastinal disorders | 1,010 | 0.6253 | 0.4564 |
| skin and subcutaneous tissue disorders | 2,082 | 0.6540 | 0.4804 |
| vascular disorders | 921 | 0.7271 | 0.5337 |
| surgical and medical procedures | 57 | 0.6789 | 0.5532 |
| social circumstances | 67 | 0.5133 | 0.4257 |

**Supplementary Table S4.** Case Study – Carbamazepine

| ADR | Real Frequency | Predicted Frequency |
|---|---|---|
| anaphylactic shock | 1 | 1.4414 |
| angioedema | 1 | 1.0333 |
| pancytopenia | 1 | 1.3838 |
| jaundice cholestatic | 2 | 1.7361 |
| hepatic enzyme increased | 3 | 2.8745 |
| eczema | 3 | 3.2447 |
| rash erythematous | 3 | 2.8145 |
| hypoglycaemia | 3 | 3.4577 |
| muscle spasms | 4 | 4.0551 |
| nasopharyngitis | 4 | 4.2663 |
| dermatitis | 4 | 3.9497 |
| dysgeusia | 4 | 4.004 |
| headache | 4 | 4.0114 |
| nausea | 5 | 4.5424 |
| diarrhoea | 5 | 4.7361 |

**Supplementary Table S5.** Case Study – Terbinafine

| ADR | Real Frequency | Predicted Frequency |
|---|---|---|
| anaphylactic shock | 1 | 1.4414 |
| angioedema | 1 | 1.0334 |
| pancytopenia | 1 | 1.3838 |
| jaundice cholestatic | 2 | 1.7361 |
| hepatic enzyme increased | 3 | 2.8745 |
| eczema | 3 | 3.2447 |
| rash erythematous | 3 | 2.8146 |
| hypoglycaemia | 3 | 3.4577 |
| muscle spasms | 4 | 4.0552 |
| nasopharyngitis | 4 | 4.2663 |
| dermatitis | 4 | 3.9498 |
| dysgeusia | 4 | 4.0041 |
| headache | 4 | 4.0115 |
| nausea | 5 | 4.5424 |
| diarrhoea | 5 | 4.7361 |

**Supplementary Table S6.** Case Study – Dyspnoea

| Drug | Real Frequency | Predicted Frequency |
|---|---|---|
| haloperidol | 1 | 1.1224 |
| trimethoprim | 1 | 1.2082 |
| amantadine | 2 | 2.3734 |
| doxepin | 2 | 2.2891 |
| paroxetine | 3 | 2.9413 |
| fondaparinux | 3 | 3.3272 |
| irbesartan | 3 | 2.8210 |
| zaleplon | 3 | 2.9550 |
| pregabalin | 4 | 3.9405 |
| dalteparin | 4 | 3.8620 |
| ziprasidone | 4 | 3.9301 |
| cabazitaxel | 4 | 4.0924 |
| cidofovir | 5 | 4.7657 |
| eltrombopag | 5 | 4.9373 |
| belinostat | 5 | 4.7895 |

**Supplementary Table S7.** Case Study – Agitation

| Drug | Real Frequency | Predicted Frequency |
|---|---|---|
| haloperidol | 1 | 1.2679 |
| rabeprazole | 2 | 2.2773 |
| sertraline | 3 | 3.3301 |
| entacapone | 3 | 3.3298 |
| nabumetone | 3 | 3.1638 |
| estazolam | 3 | 2.8091 |
| eszopiclone | 3 | 2.9520 |
| zaleplon | 3 | 2.8220 |
| lisdexamfetamine | 4 | 4.3919 |
| ziprasidone | 4 | 3.9524 |
| ziprasidone | 4 | 3.9524 |
| chlorambucil | 4 | 4.2543 |
| remifentanil | 4 | 3.9524 |
| epoprostenol | 5 | 4.5838 |
| cytarabine | 5 | 4.6928 |
